# Retinal horizontal cells use different synaptic sites for global feedforward and local feedback signaling

**DOI:** 10.1101/780031

**Authors:** Christian Behrens, Yue Zhang, Shubhash Chandra Yadav, Silke Haverkamp, Stephan Irsen, Maria M. Korympidou, Anna Schaedler, Karin Dedek, Robert G. Smith, Thomas Euler, Philipp Berens, Timm Schubert

## Abstract

In the outer plexiform layer (OPL) of the mouse retina, two types of cone photoreceptors (cones) provide input to more than a dozen types of cone bipolar cells (CBCs). This transmission is modulated by a single horizontal cell (HC) type, the only interneuron in the outer retina. Horizontal cells form feedback synapses with cones and feedforward synapses with CBCs. However, the exact computational role of HCs is still debated. Along with performing global signaling within their laterally coupled network, HCs also provide local, cone-specific feedback. Specifically, it has not been clear which synaptic structures HCs use to provide local feedback to cones and global forward signaling to CBCs.

Here, we reconstructed in a serial block-face electron microscopy volume the dendritic trees of five HCs as well as cone axon terminals and CBC dendrites to quantitatively analyze their connectivity. In addition to the fine HC dendritic tips invaginating cone axon terminals, we also identified “bulbs”, short segments of increased dendritic diameter on the primary dendrites of HCs. These bulbs are located well below the cone axon terminal base and make contact to other cells mostly identified as other HCs or CBCs. Using immunolabeling we show that HC bulbs express vesicular gamma-aminobutyric acid transporters and co-localize with GABA receptor γ2 subunits. Together, this suggests the existence of two synaptic strata in the mouse OPL, spatially separating cone-specific feedback and feedforward signaling to CBCs. A biophysics-based computational model of a HC dendritic branch supports the hypothesis that the spatial arrangement of synaptic contacts allows simultaneous local feedback and global feedforward signaling.

## Introduction

At the very first synapse of the mouse visual system, the signal from the cone photoreceptors (cones) is relayed to second-order neurons: each cone axon terminal has more than 10 output sites, contacting a sample of the 13 types of cone bipolar cells (CBCs) (Wässle et al. 2009; Tsukamoto and Omi 2014; Behrens et al. 2016), which relay the signal ‘vertically’ to retinal output neurons, the retinal ganglion cells. In a complementary circuit, laterally-organized horizontal cells (HCs) modulate the photoreceptor-BC synapse (Haverkamp, Grünert, and Wässle 2000). Each ON-cone bipolar cell (ON-CBC) dendrite invaginates the cone axon terminal to contact an individual cone output site and forms a triad with HC dendritic processes, whose distal dendritic tips also invaginate the synaptic cleft, flanking ON-CBC dendrites. In contrast, OFF-CBC dendrites contact the cone axon terminal base without forming contacts with HC dendritic tips (reviewed in Diamond 2017) (Fig. 1a).

**Figure 1.**
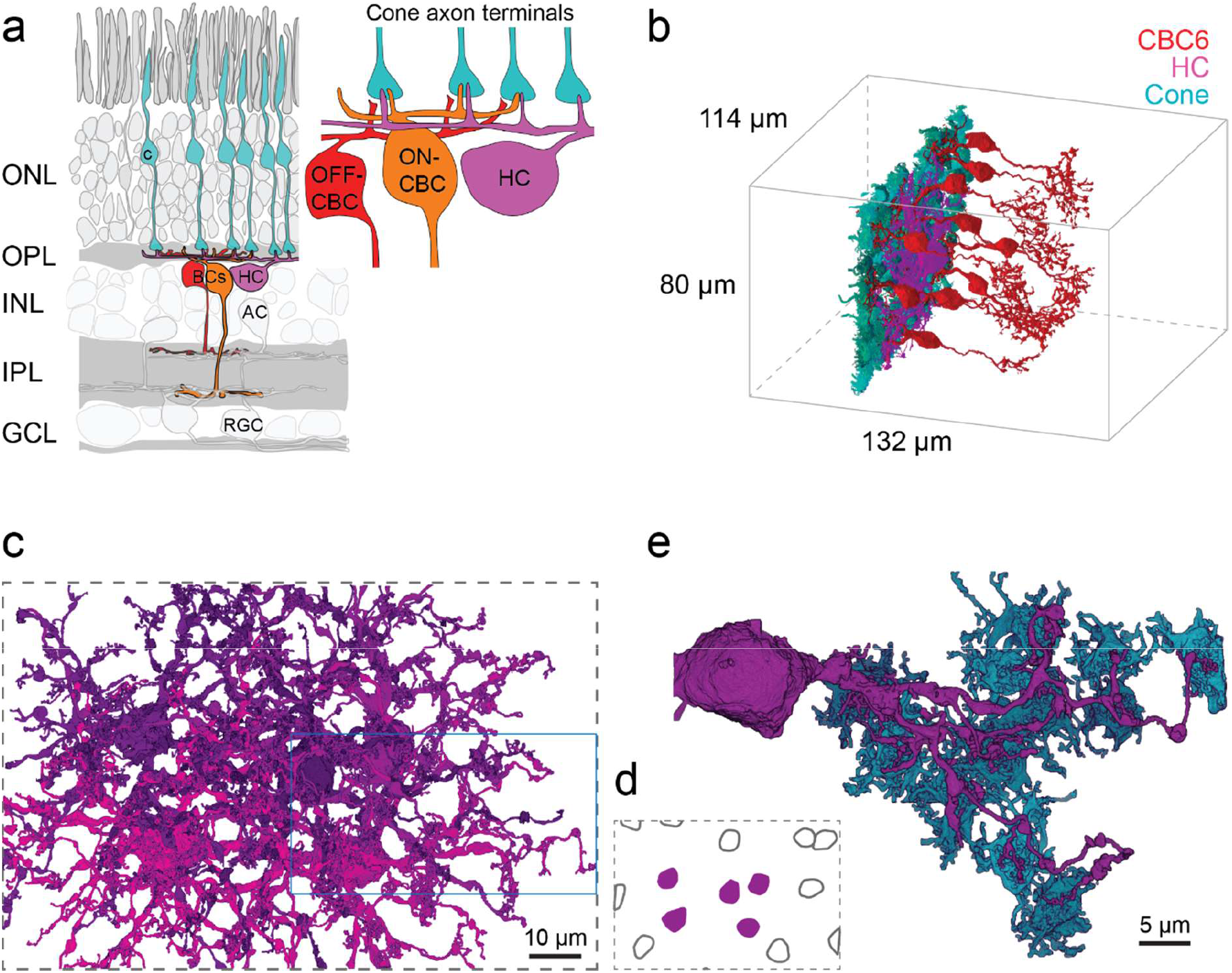
Horizontal cell reconstruction. (a) Schematic of a vertical section through the mouse retina, highlighting the reconstructed cell types. Inset: Textbook-view of the connectivity of bipolar cells (BCs) and horizontal cell (HCs) at the cone axon terminals with invaginating HC (magenta) and ON-CBC (orange) dendritic tips and basal OFF-CBC contacts (red). AC, amacrine cell; RGC, retinal ganglion cell. (b) Outlines of the dataset with volume reconstructed cone axon terminals (cyan), one HC (magenta) and several CBC6 (red, 10 of 45 CBC6s shown). (c) Volume reconstructions of five HCs (top view); blue rectangle: location of dendrite shown in (e). (d) Soma locations of the five reconstructed HCs (magenta) and 10 HCs not reconstructed (black outline) HCs. (e) Bottom view of the volume reconstruction of a complete HC dendrite (magenta) with contacted cone axon terminals (cyan). HC dendrite taken from inset in (c).

Horizontal cells are thought to play a major role in global visual processing and to contribute to contrast enhancement and generation of center-surround receptive fields, providing global feedback signals to cones and feedforward signals to BCs (reviewed in Thoreson and Mangel 2012; Drinnenberg et al. 2018; Ströh et al. 2018). Functional measurements indicate that HCs provide local feedback to photoreceptors (Jackman et al. 2011; Chapot et al. 2017), modulating each cone’s output individually. However, the role of this local feedback is unclear in the context of the HC’s traditional role of providing global feedback.

In addition, synapses between HCs and BCs were observed in in electron microscopic studies over three decades ago but have not been further investigated (Olney 1968; Kolb 1977; Linberg and Fisher 1988). Thus, despite our increasing knowledge of the complex interplay of different synaptic mechanisms underlying HC feedback to cones (Liu et al. 2013; Kemmler et al. 2014; Vroman et al. 2014; Grove et al. 2019), a quantitative anatomical picture of the outer retinal connectivity with the HC as a central player is missing.

Here, we made use of the serial block face electron microscopy dataset e2006 (Helmstaedter et al. 2013) to reconstruct the outer mouse retina with a focus on the HC circuitry and identify the connectivity motifs made between HCs and other neuron types. In addition to the invaginating contacts between cones and HCs in the cone axon terminal, we identified and quantitatively assessed – at the level of primary HC dendrites – putative GABAergic synapses among HCs as well as between HCs and CBCs as previously hypothesized (Dowling, Brown, and Major 1966; Marchiafava 1978; Yang and Wu 1991; Duebel et al. 2006). Based on a biophysical model of HC signaling, we propose that a role of this putative second synaptic site in HCs is to provide global signals in the form of GABAergic input to postsynaptic CBCs, complementing the local feedback provided directly to cones. This suggests that a single interneuron can simultaneously provide local reciprocal feedback and global feedforward signals at distinct synaptic locations.

## Results

### Reconstruction of horizontal cells and connectivity with cone photoreceptors

Using the publicly available serial block-face electron microscopy dataset e2006 (Helmstaedter et al., 2013, Fig. 1b), we reconstructed five complete dendritic arbors of HCs in the outer mouse retina (Fig. 1c,d). We analyzed the contacts of all three classes of neurons in the outer retina – cones, BCs and HCs (Helmstaedter et al. 2013; Behrens et al. 2016) – to gain a complete picture of outer retinal connectivity. The reconstructed HCs had a dendritic area of 4,600 ± 400 μm^2^ (mean ± SD) with 4 to 6 primary dendrites leaving the soma (n = 5 HCs) and extended fine dendritic tips towards cone axon terminals (Fig. 1e). Next, we analyzed the HC connectivity within the cone axon terminals (Fig. 2a). Each HC contacted on average 61 ± 5 (between 51 and 77) cones; these were all cones within its dendritic field. Interestingly, we found that cones closer to the HC somata were contacted with significantly more fine HC tips than distal ones (Fig. 2b,c; Generalized Additive Model with smooth term for distance from soma and random effect of specific HC, *p* < 2 · 10^−16^ for smooth term, see Methods for details). In addition, the contact area – the area of close apposition between cell membranes, a proxy for the probability of synaptic contacts (Helmstaedter et al. 2013) – between an individual HC and cones followed the number of invaginating contact sites along the HC dendrite and significantly decreased towards the HC’s periphery (Fig. 2d; Generalized Additive Model with smooth term for distance from soma and random effect of specific HC, *p* = 4.2 · 10^−7^ for smooth term). Together this suggests that HCs get most input from cones close to their soma.

**Figure 2.**
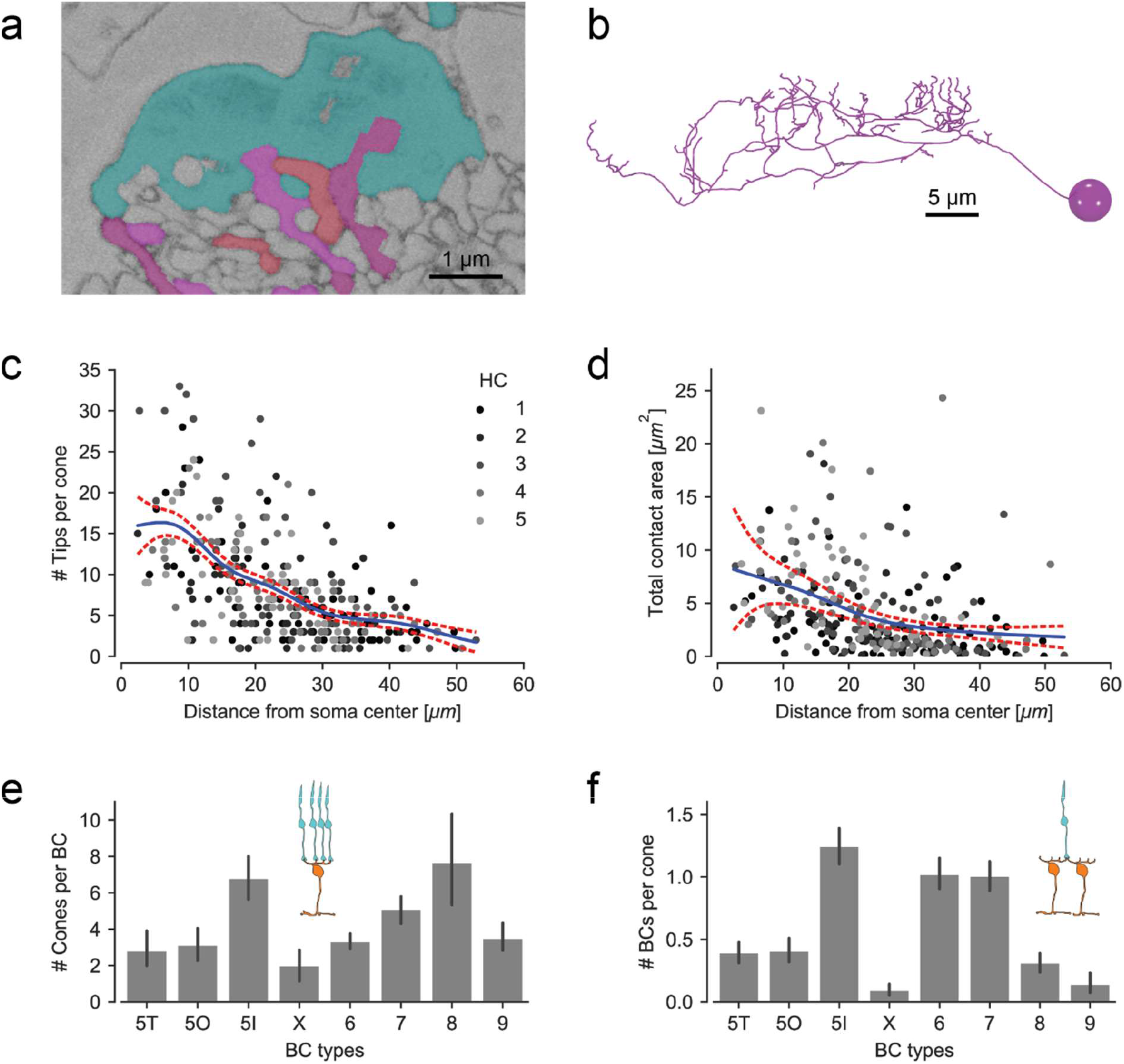
Horizontal cell-to-cone and HC-to-ON-CBC contacts. (a) EM slice showing a cone axon terminal (cyan) with invaginating contacts from an ON-CBC (red) and a HC (magenta). (b) Vertical view of the skeleton model of the HC branch from Fig. 1e illustrating the increase of the number of dendritic tips towards the soma. (c) HC skeleton tips per contacted cone vs. distance from HC soma. Blue: Poisson GAM fit with 95%-confidence interval (red). (d) Contact area between HC and cone axon terminal volume reconstructions per cone vs. distance from HC soma. Blue: Gamma GAM fit with 95%-confidence interval (red). (e) Contacted cones per BC for all CBC types. (f) BCs contacted per cone for all CBC types. Number of ON-BCs contacting each cone per type. Both (e) and (f) redrawn using data from Behrens et al. 2016. Error bars show 95% CI.

### Invaginating contacts between cones, ON-cone bipolar cells and horizontal cells

We found that the connectivity between cones and their postsynaptic partners was the typical “triad synapse” motif, where HC tips in the synaptic cleft are closely associated with invaginating dendrites of ON-CBCs. A previous study found that some ON-CBC types such as CBC types 5T, 5O, 8 and X sampled sparsely from cones (Fig. 2e; see also Fig. 3 in Behrens et al. 2016). In addition, the CBCX made rather small basal but not ‘typical’ invaginating contacts at the cone axon terminal, more resembling OFF-CBC contacts. If these BC types contacted cones more sparsely, the number of contacts with invaginating HC dendrites should be lower as well. We checked all ON-CBC contacts (n = 36) at five central cones (making contacts with all 5 reconstructed HCs) and identified one or two invaginating HC dendritic tips per ON-CBC tip for all contacts. This typical “triad” motif implies that the number of contacts between HCs and ON-CBCs matches the number of cone contacts per CBC. Thus, the number of contacts between HCs and BC types CBC5T, 5O, 8 and X within the cone axon terminal is lower than for the other ON-CBC types (Fig. 2f).

**Figure 3.**
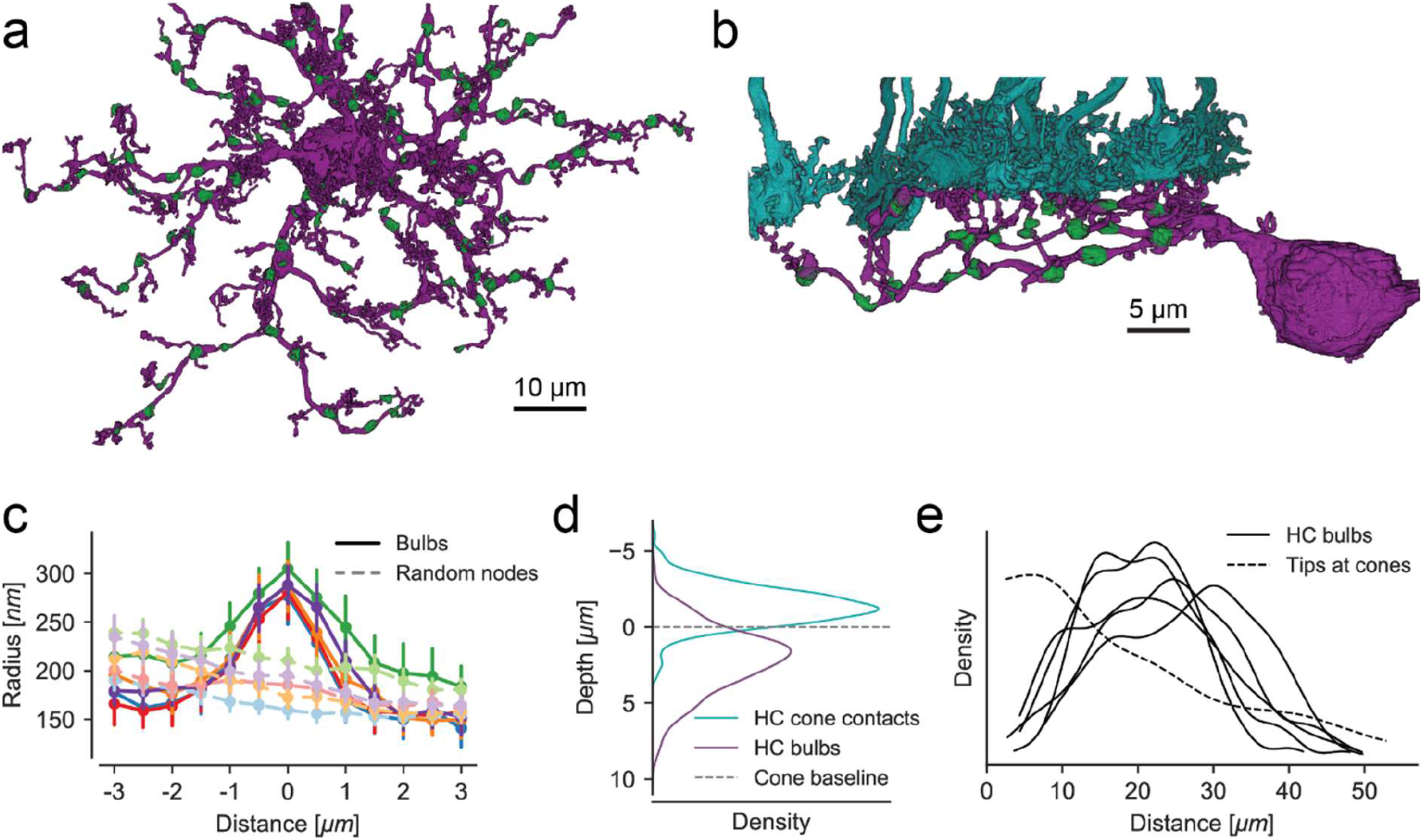
Identification of bulbs in horizontal cells. (a) Top view of a reconstructed HC with bulbs highlighted in green. (b) Side view of a branch from the same HC with bulbs (green) and cone axon terminals (cyan). (c) Dendritic radius profile at bulb locations (solid curves w/high saturation) compared to randomized points on the dendrites (dashed curves w/low saturation) with matching distribution of distances from soma and tips. (d) Depth of bulbs compared to HC-to-cone contacts. (e) Kernel density estimate of the distance distribution of bulbs relative to the soma for all five HCs. Dashed line: Model fit showing distribution of HC skeleton tips at cones from Fig. 2c.

### Non-invaginating (bulb) contacts between horizontal cells and bipolar cells

Horizontal cell feedback modulates the cones’ glutamate release directly within the invaginating cleft (Kamermans et al. 2001). For a spatially uniform stimulus pattern, this feedback should be spatially correlated and should affect the output of all cones similarly. However, for a spatially uncorrelated stimulus pattern, HC distal tips would receive uncorrelated input and provide uncorrelated feedback to neighboring cones. Indeed, HC dendritic tips have been shown to feature highly localized signals (Jackman et al. 2011; Chapot et al. 2017), making it possible to relay highly localized feedback to individual cones. Yet, in addition to transmitting local uncorrelated signals, for spatially uniform stimuli, this pathway may be also responsible for global signals traditionally suggested as a general role of HCs (Thoreson and Mangel 2012; Drinnenberg et al. 2018). The reason is that, while the local feedback from a HC distal tip to its presynaptic cone will attenuate the local signal received by the HC distal tip, the global signal transmitted from HC dendrites and soma is not attenuated.

However, a second synaptic output pathway where HCs could provide direct GABAergic output to BC dendrites independent from the invaginating cleft has been postulated (Marchiafava 1978; Yang and Wu 1991; Duebel et al. 2006). Because dendrites of OFF-CBCs have a low and dendrites of ON-CBCs have a high chloride level (Duebel et al. 2006), GABA released by HCs that binds to GABA-gated chloride membrane channels in CBC dendrites will hyperpolarize an OFF-CBC and depolarize an ON-CBC. Therefore, an antagonistic surround in both ON- and OFF-CBCs can be generated by HCs through this lateral feedforward pathway. This feedforward signaling pathway could integrate both correlated and/or uncorrelated HCs signals along the HC primary dendrites generating a global output signal which is then relayed to BCs. However, such a synaptic HC-BC connection has not been identified so far.

In search of this synaptic site between HCs and CBCs, we systematically examined the five volume-rendered HCs and found regularly distributed, dendritic swellings along the primary dendrites (Fig. 3a,b). These dendritic swellings (bulbs) showed a marked increase in dendritic diameter (Fig. 3c). Almost all bulbs were located clearly below the cone axon terminal base and not in direct contact with it (Fig. 3b,d). In contrast to invaginating dendritic tips that showed a higher density towards the soma of the HC, the bulbs were more regularly distributed along the primary dendrites (Fig. 3e).

Most of the identified bulbs contacted either bulbs of other volume-rendered HCs (67 out of 545, Fig. 4a,c) or dendrites of ON- and OFF-CBCs (219 out of 545; Fig. 4b,d) or both (69), suggesting that the bulb structures represent potential HC-HC and/or HC-BC synapses. For the remaining 190 bulbs, we had no information about the identity of the contacted cells. As only five HCs were traced, these contacts may well represent contacts to other HCs and/or BCs that were not traced because their soma was located outside the EM stack.

**Figure 4.**
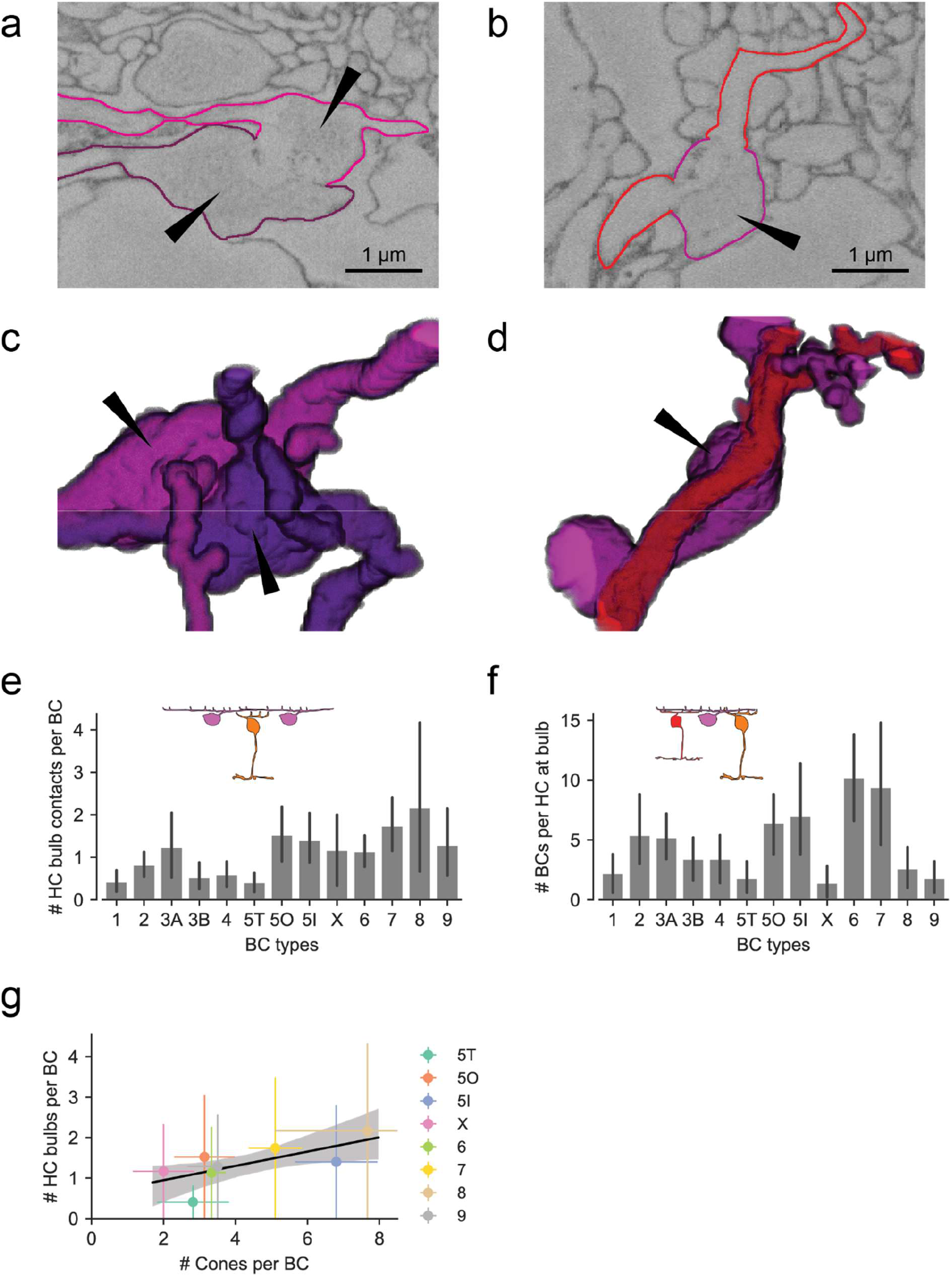
Horizontal cell bulb contacts with other neurons. (a) EM slice showing bulb contact between two HCs. (b) Bulb contact between HC (magenta) and ON-CBC (red). (c) Volume-rendered contact site between bulbs from two HCs. (d) Volume-rendered contact site between HC (magenta) bulb (arrowhead) and ON-CBC dendrite (red). (e) HC bulb contacts per BC for all CBC types. (f) BCs contacted by bulbs per HC for all CBC types. (g) Bulb contacts vs. contacted cones for all ON-CBC types (data from (e) and Fig. 2e) with linear regression. All error bars show 95% CIs.

Interestingly, we found a difference in bulb-level connectivity between HCs and OFF-CBCs vs. HCs and ON-CBCs: the majority of ON-CBCs contacted HCs at the bulb site (except for CBC5T); all OFF-CBC types made considerably fewer contacts than ON-CBCs (except CBC3A) (Fig. 4e). However, the overall number of contacts per CBC is likely underestimated since contacts to not reconstructed HCs are not included (see above). Furthermore, the number of BCs contacted at bulbs per HC was lowest for the CBC types 5T, X, 8 and 9 and highest for CBC types 6 and 7 (Fig. 4f). For types X, 8 and 9, the low numbers likely originate in their lower cell count while for CBC5T (which has the same dendritic density as CBC5O and 5I cells) it is a consequence of the low number of contacts per CBC. Comparing the bulb-to-ON-CBC contacts with the number of cone-to-ON-CBC contacts taken from our recent study (Behrens et al. 2016) showed that both connectivity patterns are almost identical for nearly all BC types (Fig. 2e,4e). The only striking difference was found within the group of CBC5 cells: CBC5T has the lowest contact number with both HC bulbs and cones whereas CBC5I made many contacts with both cones and bulbs. CBC5O sampled from as few cones as CBC5T but made more bulb contacts, similar to CBC5I (Fig. 4g). Thus, while the morphological properties such as density, dendritic field size, axon terminal size and stratification depth of the three ‘sister’ types of CBC5 do not differ much, they can be distinguished based on their connectivity patterns with cones and HCs in the outer retina.

### Horizontal cell bulbs are putative synaptic structures

If HC bulbs indeed formed presynaptic sites to relay global cone signals, one would expect presynaptic proteins to be co-localized with these bulbs. In mammalian HCs, several presynaptic proteins are associated with synaptic vesicles, including the vesicular gamma-aminobutyric acid transporter (VGAT) (Guo et al. 2009; Cueva et al. 2002). However, most immunolabeling studies of the OPL have focused on the distal tips of HCs rather than their primary dendrites. To identify potential presynaptic sites along HC dendrites, we therefore used calbindin and VGAT antibodies to visualize HCs and presynaptic vesicles, respectively (Fig. 5a). Indeed, we found intense VGAT staining in dendritic thickenings at the same depth at which bulbs were found in the EM data.

**Figure 5.**
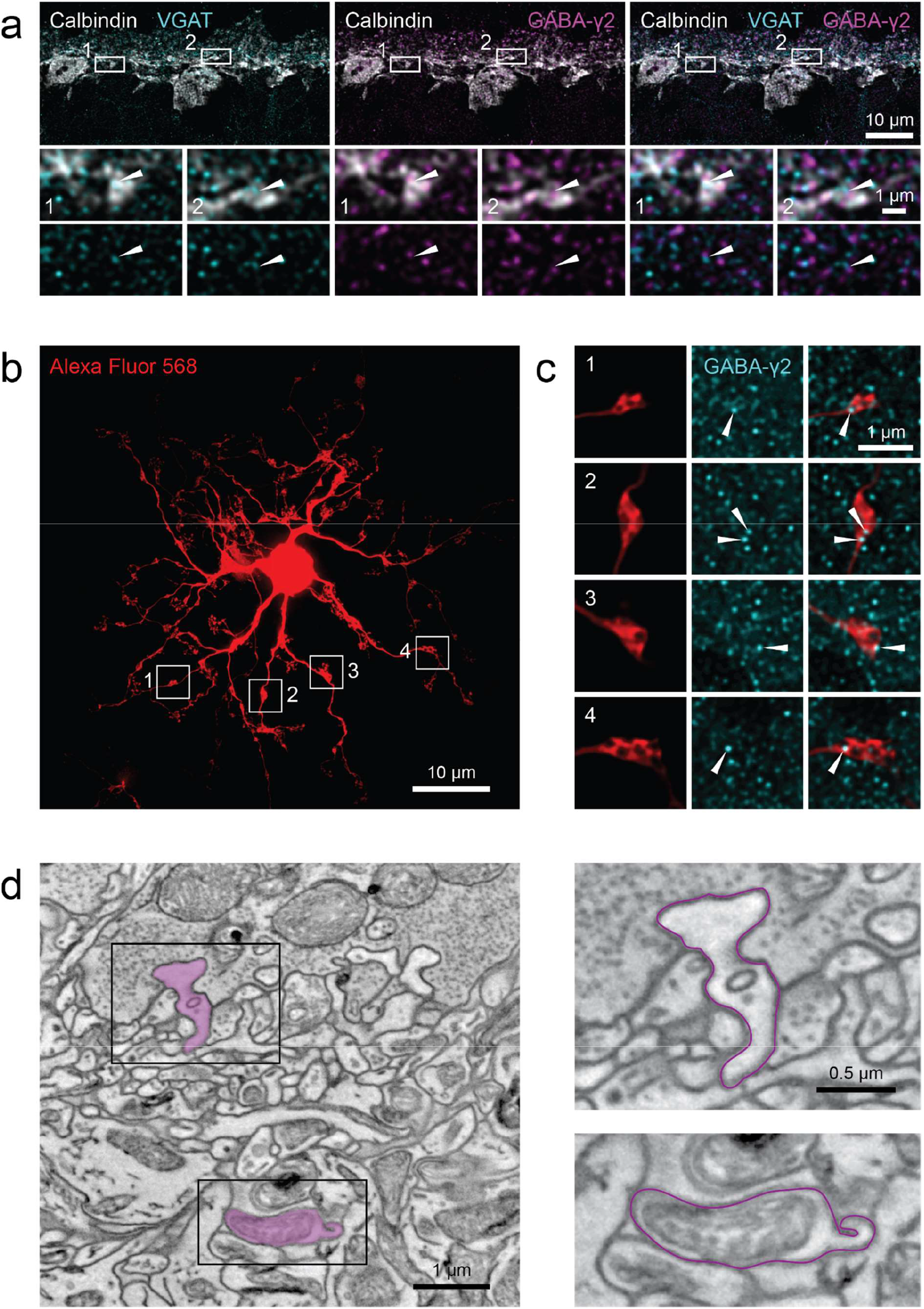
Synaptic structures at horizontal cell bulb contacts. (a) Calbindin labeled HCs with VGAT (cyan) and GABA receptor gamma2 (magenta) immunolabeling in vertical outer retinal section. Arrowheads indicate co-localization on primary dendrites. (b) Alexa Fluor 568-injected HC with identified bulbs (white boxes indicate examples). (c) Enlarged bulbs (red) from boxes in (b) with GABA receptor γ2 immunolabeling (cyan). Arrowheads indicate co-localization. (d) Electron microscopy image showing a manually traced HC (magenta) with a dendritic tip invaginating in the cone axon terminal (upper right) and the primary dendrite below the cone axon terminal with a HC bulb of the same cell (lower right). Note the mitochondrial structure in the bulb. Black rectangles: location of magnifications shown on the right.

If bulbs are the site of GABAergic synapses between HCs and to BCs, then GABA receptors should be present at VGAT-positive bulbs as well. In the mouse retina, different GABA receptor subunits are expressed in the outer retina: A ‘dashed’ band of a GABA receptor α1 subunit staining can be seen at the level of the cone axon terminals (Haverkamp and Wässle 2000), indicating that α1 subunits are prominently expressed by HC dendritic tips invaginating in the synaptic cleft (Kemmler et al. 2014). In contrast, GABA receptor γ2 subunits have a broader expression profile that clearly stratifies below the cone axon terminals (Haverkamp and Wässle 2000). Indeed, we found that VGAT and GABA receptor γ2 subunit immunolabeling co-localized on bulb-like structures (representative example shown in Fig. 5a; similar results were obtained in four out of four immunostainings from two different mice). To assess GABA receptor distribution on individual bulbs, we injected HCs in the whole-mount preparation with the fluorescent dye Alexa Fluor 568 and then performed GABA receptor immunolabeling (Fig. 5b,c). The γ2 subunit immunoreactivity was strong at the level of the primary dendrites and all identified bulbs (n = 30 bulbs in n = 3 injected HCs) showed immunolabeling for the GABA receptor γ2 subunit, indicating that bulbs may provide and/or receive GABAergic input (Fig. 5c). Sources for GABAergic input may be other HCs (Liu et al. 2013) or eventually interplexiform amacrine cells (Witkovsky, Gábriel, and Križaj 2008; Dedek et al. 2009).

For further evidence that the GABA receptors in bulbs may be synaptic structures, we performed focused ion beam scanning EM and reconstructed HCs from their dendritic tips in the invaginating cleft (Fig. 5d, left) to the depth in the OPL where bulbs are located. In this EM image stack, bulbs could be identified based on their thickened structure (Fig. 5d, supplementary image stack). These structures always contained mitochondria which are typically found in presynaptic structures (n = 7 bulbs) (Gala et al. 2017). However, compared with the glutamate-filled vesicles in the cone axon terminal, the vesicles in both invaginating tips and bulbs of HCs were barely detectable (Fig. 5d). Thus, the vesicle distribution was not further investigated (see Discussion).

### Biophysical modelling indicates potential bulb function

To study the potential functional role of HC bulb contacts, we built a biophysical model of a HC dendritic branch with cone input (Fig. 6a,b) based on our previously published model (Chapot et al. 2017). We stimulated the cones in the model with either full-field or checkerboard noise for spatially correlated and uncorrelated input, respectively, and measured voltage signals in the HC dendritic tips invaginating into cone axon terminals (n = 12) and in the bulb structures (n = 14) (Fig. 6c,d). For full-field stimuli, we found high correlations between voltage signals from all recording points (0.85 ± 0.10). Due to vesicle release noise included in the model, which occurred independently at each synapse between cones and HC tips, correlations between signals in different tips (0.73 ± 0.07) and between signals in tips and bulbs (0.84 ± 0.06) were lower than those between signals in bulbs (0.94 ± 0.04) (Fig. 6e). However, correlations between bulbs and the average over the tip signals (0.97 ± 0.02) were similar to the correlations between bulbs.

**Figure 6.**
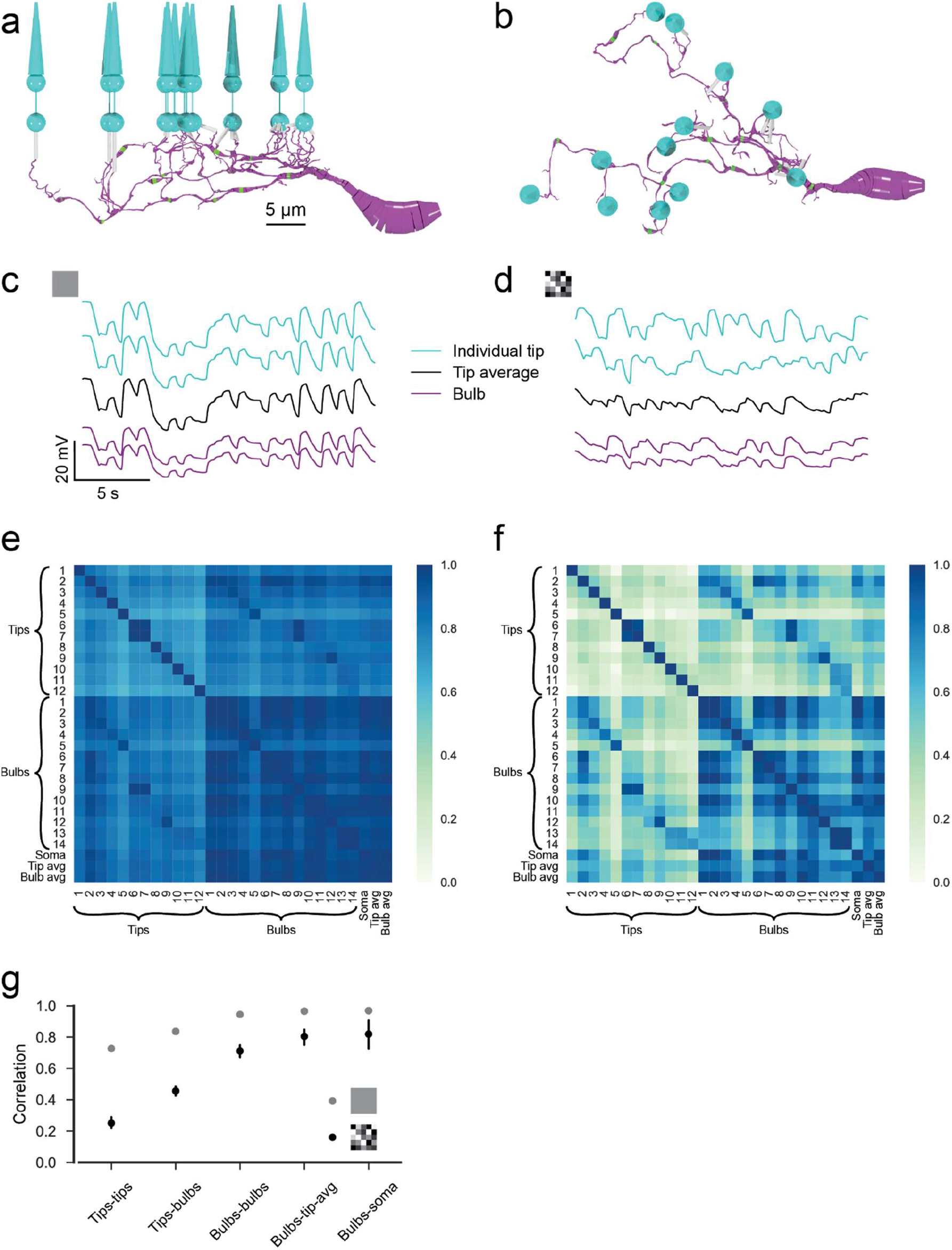
Biophysical modelling. (a) Side and (b) top view of the modelled HC dendrite with bulbs (green) and cones (cyan). (c) & (d) Example voltage traces recorded at HC dendritic tips below cones, at bulbs and average over all tips for (c) a random full-field noise stimulus and (d) a random checkerboard noise stimulus without synaptic vesicle release noise. (e) & (f) Correlations from 60 s of (e) full-field and (f) checkerboard stimulation including synaptic vesicle release noise. (g) Mean correlations between different tips, between tips and bulbs, between different bulbs, bulbs and the tip mean and bulbs and the soma for both stimuli. avg, average.

For the checkerboard noise, which is a spatially maximally uncorrelated stimulus, the result was different: The average correlation between voltage signals in tips was rather low (0.25 ± 0.14), confirming our previous results (Chapot et al. 2017). In contrast, the average correlation between voltage signals in bulbs was much higher (0.71 ± 0.19) and so was the correlation between bulb signal and the average over the tip signals (0.80 ± 0.10) (Fig. 6g). Together, this indicates that for a natural stimulus with spatially uncorrelated stimulation pattern, the global component of the stimulus dominates the signal at the level of the bulbs whereas the local signal at the HC dendritic tips can be used for feedback to individual cones (Fig. 7).

**Figure 7.**
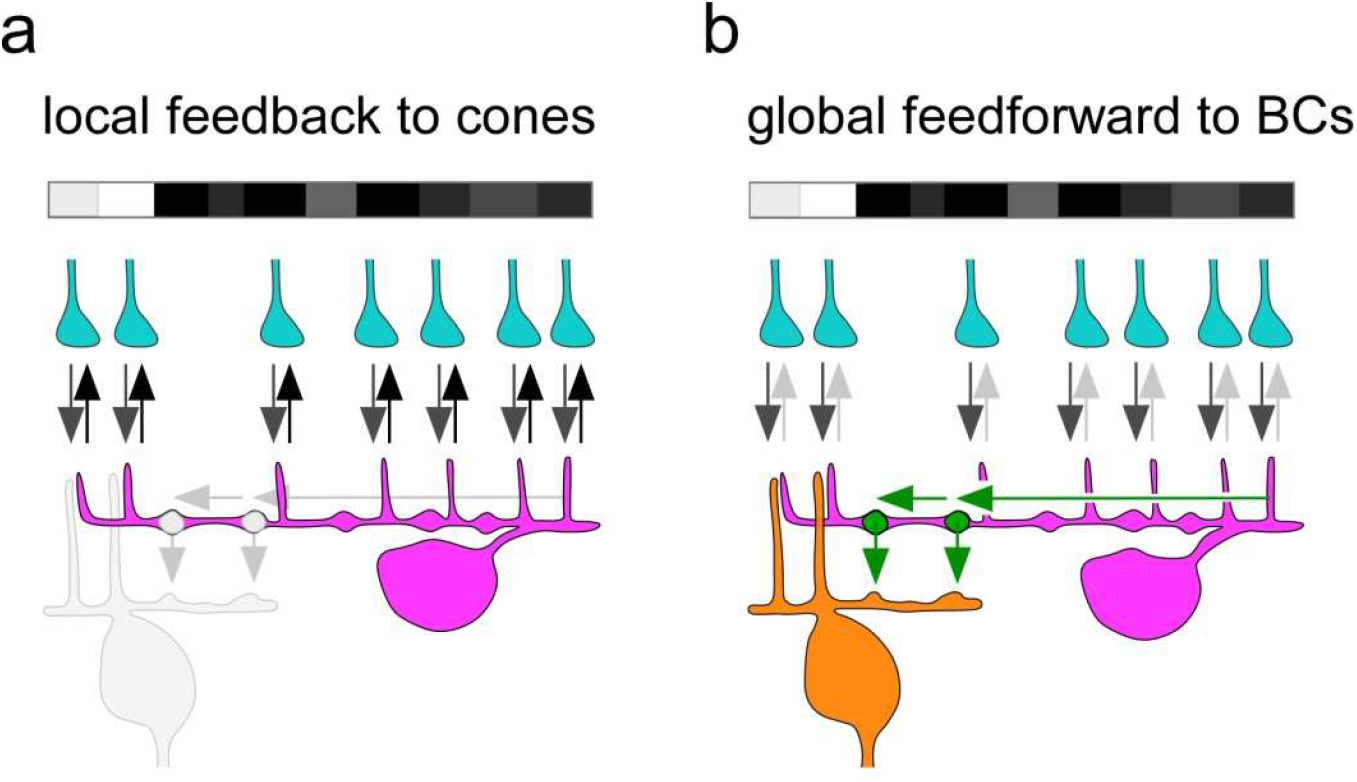
Synaptic circuitry of horizontal cells in the outer mouse retina. (a) Dendritic tips of a HC (magenta) receive cone input (grey arrows) and provide local and cone-specific feedback (black arrows) to cone axon terminals (cyan) in the presence of a spatially and temporally uncorrelated white noise stimulus (white/grey/black bar). (b) For the same uncorrelated stimulus, the cone input signals (grey arrows) are integrated to a global signal in the HC dendrite (green arrow) and forwarded by a BC-contacting bulb (green) to the BC (orange) forming the surround signal.

## Discussion

Classically, HCs have been postulated to perform global computations like gain control and contrast normalization (Barnes, Merchant, and Mahmud 1993; Verweij, Kamermans, and Spekreijse 1996; Drinnenberg et al. 2018). However, local signaling in HCs has recently been reported in a number of studies (Jackman et al. 2011; Chapot et al. 2017; Grove et al. 2019), raising the important question, how a strongly electrically coupled syncytium of a laterally organized interneuron can perform both local and global computations and relay them to its presynaptic partners. Here, we provide a quantitative picture of connectivity in the outer retina including HCs and describe a new putative synaptic site in HCs. In contrast to the well-described feedback synapse that modulates the cone output, this putative synapse is likely a feedforward synapse that provides GABAergic drive to other HCs as well as to BCs. On the functional level, it likely relays an integrated, global signal, resulting from multiple local signals from numerous cones. This global output signal may contribute to the global center-surround organization of bipolar cells, as has long been posited (Werblin and Dowling 1969; Schwartz 1974). (Fig. 7).

### Horizontal cell bulbs are likely synaptic structures

Based on few ultrastructural electron microscopic examples of chemical synapses between HCs and BCs shown in earlier years (Olney 1968; Kolb 1977; Linberg and Fisher 1988), these direct synaptic connections have long been suggested (Miller and Dacheux 1976; Marchiafava 1978; Yang and Wu 1991; Vardi et al. 2000; Duebel et al. 2006), However, these connections have never been systematically investigated. Here, we used the e2006 EM data set that was initially used to describe inner retinal connections (Helmstaedter et al. 2013) and photoreceptor-to-BC synapses (Behrens et al. 2016). The majority (~65%) of identified HC bulbs was contacted by other HC bulbs and/or BC dendrites indicating that bulbs were not randomly distributed dendritic thickenings. This finding was supported by the uniform distribution of bulbs across the HC dendritic tree suggesting the overall mosaic organization of the mouse retina is maintained at this synaptic level. Furthermore, we found that the bulbs contained mitochondria required to supply energy for the presynaptic vesicle release mechanism as described for a reciprocal amacrine cell synapse in the cat retina (Ellias and Stevens 1980). Indeed, we found synaptic proteins on HC bulbs such as VGAT and GABA γ2 receptors on HC bulbs that represent pre- and postsynaptic markers, respectively. However, which retinal cell type(s) express the GABA receptors is still unclear. Since the bulbs contact other HC bulbs as well as BC dendrites, it is conceivable that the bulbs represent a general GABAergic output site to postsynaptic cells (reviewed in Diamond 2017).

Unfortunately, we could not detect any neurotransmitter vesicles at HC bulbs in our EM images – a problem that was reported for HCs decades ago (Dowling and Boycott 1966) and may be attributed to methodological limitations of the dataset. However, in an earlier cat retina study, vesicle clusters in HC dendrites were found to be “…typically organized around some denser particles…” (Fig. 18 in Kolb 1977). In addition, in the mouse retina, “.the synaptic complex, was of conventional configuration, involved the horizontal cell presynaptically, and was always found in the innermost aspect of the outer plexiform layer.” (Fig. 11 in Olney 1968) – this is where we found the majority of bulbs in our study. Another reason why we did not find vesicles could be the (somewhat unlikely) GABA secretion by GABA transporters as shown in the fish retina (Schwartz 1987). Additionally, we also cannot exclude the possibility that bulbs receive direct glutamatergic input by diffusion from photoreceptors – similar to OFF-CBCs – however, the distance of HC bulbs from the release sites in cones axon terminals appears to be relatively large compared with the OFF-CBC dendrites that directly contact the axon terminal. Hence, we propose that GABA release at the bulbs results from electrical signal propagation along the dendrites of the HCs rather than being initiated via diffusing glutamate.

A GABAergic synapse between HCs along with a depolarizing chloride level in HCs (Miller and Dacheux 1976; Kamermans and Werblin 1992) could transmit a lateral signal between them, similar to the lateral signal spread within the syncytium formed by electrical synapses. One might therefore imagine that mouse HC bulbs could represent sites of gap junctions - electrical synapses between HCs (He, Weiler, and Vaney 2000). However, the vast majority of connexin57 protein forming electrical synapses in mouse HCs is expressed at more distal sites on the HC dendrites and is not very prominent on the bulb-bearing primary dendrites (Janssen-Bienhold et al. 2009). Thus, it is unlikely that the role of HC bulbs is to couple HCs by electrical synapses.

Our finding of an additional synapse in the outer mouse retina may explain how HCs may perform both feedback to cones and feedforward signaling to BCs simultaneously, which may contribute to understanding a longstanding enigma.

### Selective connectivity with ON-CBC types as a mechanism of synaptic scaling?

As previously reported, we found some ON-CBCs contact cone axon terminals in a very specific manner. For example, the CBC types 5T, 5O, X, 8 and 9 contact considerably fewer cones than expected from their relatively large dendritic field whereas other types such as types 5I, 6 and 7 contact almost every cone located within their dendritic field (Behrens et al. 2016). Remarkably, this connectivity is also reflected in the number of bulbs connected by ON-CBCs: Types 5T, X and 9 contact fewer bulbs than the other types while type 8 contacts only slightly more bulbs than other cells despite its significantly larger dendritic field. This correlation of excitatory and inhibitory synapse number may be a form of synaptic scaling (Turrigiano 2011) that could have an effect on the functional organization of the receptive field of BCs. The center of a receptive field is defined as the region that is driven by excitatory (i.e., direct glutamatergic) input from cones whereas the surround is formed by the lateral inhibition by interneurons in the periphery. A balanced synaptic weight between center and surround activation is likely to be crucial for the BC’s ability to stay within the operational range of its output synapses.

The only exception is the BC type 5O; it contacts only a few cones but has relatively many bulb contacts, in strong contrast to the types 5T or 5I, which make few or many contacts to both cones and HC bulbs, respectively. Based on their morphology and their stratification depth in the inner plexiform layer, these three BC types are hardly distinguishable. However, they differ in their connectivity with cones and HC bulbs in the outer retina, and thus, may be functionally distinct regarding their receptive field properties (Franke et al. 2017). Whether the size and efficiency of synaptic contacts is different and whether or how synaptic scaling is implemented for ON-CBC and HC contacts has to be addressed in a future functional study.

### Functional consequences of a second synaptic layer in the outer retina

For decades, an open question has remained about how lateral inhibition essential for center-surround organization in the outer retina can be generated by fine structures such as the HC dendritic tips invaginating into the cone axon terminal (Yang and Wu 1991). With the new putative synaptic site described here between HCs and BCs, we propose a potential solution for this long existing dilemma of how HCs provide global/local feedback to cones and global feedforward signaling to BCs. In our view, HCs are a retinal interneuron with two functional specializations: feedback to cones and feedforward synapses to BCs and other HCs.

A recent study showed that, based on their thin diameter and high resistance, the fine HC dendritic tips are optimized for generation of local cone-specific feedback and to a lesser extent for lateral propagation of electrical signals (Chapot et al. 2017). However, the extent to which the feedback to cones is global or local strongly depends on the presented visual stimulus. For small-scale non-correlated visual stimuli, the feedback is expected to be cone-specific and can differ significantly for neighboring cones, strongly following the cone output signal. For a highly correlated visual stimulus, the feedback would be similar at all dendritic tip synapses, thus generating a rather uniform, global feedback pattern generated by correlated local activity.

The second synapse type made by HCs, the GABAergic bulb synapse, may be optimized for the relay of global signals to other HCs and BCs that are modulated in at least three different ways: First, all the photoreceptor input that reaches the primary HC dendrites and the bulb synapses would be filtered by the feedback at the dendritic tip feedback synapses. Second, the spread along HC dendrites is subject to passive filtering. Third, depending on the coupling state of the HC network, via electrical synapses or depolarizing GABAergic synapses, signals in HC dendrites may be strengthened or weakened. In any case, the global signal would be relayed to postsynaptic cells and contribute to center-surround antagonistic receptive field organization.

### Interaction between global and local signaling pathways

Under conditions with a rather uniform stimulation pattern, the local feedback from HCs to photoreceptors is thought to include global (lateral) signals (Warren et al. 2016) because of the high degree of correlation in the stimulus. Thus, HC feedback can accomplish both very local and spatially extended (i.e., global) feedback control of synaptic release. With the uniform stimulation pattern used in our model, the feedback to cones is correlated and is expected to be weak (Smith 1995). The reason is that the local dendritic tip signal in HCs itself is attenuated by the local feedback and hardly propagates over long distances along primary dendrites into other HC fine dendritic tips. Feedback is necessary to regulate the synaptic release of cones to maintain the optimal operating point in the output synapse, and thus to preserve the S/N ratio (Borghuis, Ratliff, and Smith 2018). However, strong feedback can become unstable for high feedback loop gains and typical synaptic delays (Smith 1995). To prevent instability, the HC feedback includes an ephaptic mechanism which is fast with minimal delay (Vroman et al. 2014; Chapot et al. 2017). However, the feedback strength is also limited by the need to maintain high gain at the cone output to CBCs. In contrast, ‘simple’ feedforward inhibition is always stable and can use high signaling amplitudes. These functional requirements constrain the feedback and feedforward synapses to be at structurally different synaptic sites. In addition, the synaptic mechanisms – ephaptic/pH-mediated feedback and GABAergic feedforward signaling – may contribute to the very specialized function of the two synapses: Whereas the feedback synapse can decrease but also increase the cone output signal depending on the stimulus context (Smith 1995; Kemmler et al. 2014), the bulb synapse only provides GABAergic drive. Its feedforward signal to CBCs has a different role, of providing inhibition to balance the excitation in CBCs, along with a stronger spatial surround than can be provided by the cone signal. Yet, both feedback and feedforward HC signals will contribute to the surround in CBCs and all downstream neurons.

### The horizontal cell – an interneuron with multiple functions

Interneurons show a remarkable heterogeneity and diversity in both function and morphology in all parts of the brain (reviewed in Cardin 2018). The morphology of an interneuron is thought to reflect its function: In the retina, for instance, BCs relay signals from the outer to the inner synaptic layer, whereas wide-field amacrine cells relay information laterally across the retina (reviewed in Euler et al. 2014; Diamond 2017). However, morphology can be deceiving; for example, the very symmetric starburst amacrine cells in the mammalian retina compute the direction of image motion (Taylor and Smith 2012). Moreover, some interneurons have been shown to serve more than one functional role but do so in a context-dependent manner. For example, in low light conditions, AII amacrine cells are central to the primary rod vision pathway, while under photopic conditions they change their role and contribute to approach sensitivity (Münch et al. 2009). For A17 amacrine cells (Hartveit 1999) which make reciprocal synapses with rod BCs a functional switch between local and more global processing has been suggested (Schubert and Euler 2010). More intriguing is our finding that different synaptic functions such as feedforward and feedback signaling of HCs are apparently performed simultaneously at different synaptic sites as previously shown for the VG3 amacrine cell type in the mouse retina that provides excitatory and inhibitory drive at distinct synaptic sites (Lee et al. 2016; Tien, Kim, and Kerschensteiner 2016).

An interesting question is why local and global signaling – two different synaptic tasks – are performed in the same interneuron in the mouse outer retina. Circuits of the inner retina seem rather to be built from several types of neurons, each contributing a specific function. Maybe with few exceptions – as illustrated by A17 and VG3 amacrine cells discussed above – amacrine cells, which with ~45 types are the most diverse retinal interneuron, are thought to represent specific computational tasks. Why is this motif not implemented in the outer retina? Two possible explanations may play a role here: First, the cone axon terminal system is among the most complex synaptic structures in the brain (Haverkamp, Grünert, and Wässle 2000). Therefore, integrating a second, dedicated interneuron type during evolution may have been avoided for the sake of space limitation and circuitry simplification. This hypothesis is supported by the fact that the reciprocal feedback synapses to rod photoreceptors are not provided by an additional interneuron type but by an additional intraretinal axon terminal system of HCs which is a unique structure for interneurons in the brain. Second, the cones require a global feedback signal that represents the average background, so their synaptic vesicle release can optimally represent a contrast signal (Srinivasan, Laughlin, and Dubs 1982). The most straightforward mechanism to generate such a global feedback signal is through summation of many local signals from individual cones. Although global feedback might be arranged at a separate synapse from local feedback, it is most straightforward to provide this integrated signal as local/global feedback to each cone to control its release rate.

## Methods

### Dataset

Our analysis is based on the SBEM dataset e2006 (Helmstaedter et al. 2013, https://www.neuro.mpg.de/connectomics). The dataset covers a piece of mouse retina of 80 x 114 x 132 μm with a resolution of 25 x 16.5 x 16.5 nm. We identified the somata of 15 HCs and skeletonized the dendrites of the five central HC in KNOSSOS (Helmstaedter, Briggman, and Denk 2011, www.knossostool.org). We used algorithms published with the dataset to reconstruct the volumes of HCs, BCs and cone axon terminals in the OPL and to identify their contacts (for details see Behrens et al. 2016).

We manually identified HC bulbs and their contacts. To compare the dendritic diameter profile around the bulbs with the one of random points on the dendrite (Fig. 3c), we used the Vaa3D-Neuron2 auto-tracing (Xiao and Peng 2013) to get a simplified representation of the HC morphologies from the volume reconstruction, consisting of a regularly spaced grid of nodes with associated diameters. For each bulb position we identified the closest node and extracted the dendritic diameter profile around it. For a fair comparison to average points on the dendrite, we draw a random set of nodes with distributions of average distances from soma and tips matching the bulb locations.

To calculate the statistics of HC-to-BC contacts at bulbs, we included only BCs where the center of the BC dendritic field was within the dendritic field of at least one of the reconstructed HC. With this method, the numbers in Fig. 4e,f are a lower bound. For the HCs, additional contacts on branches ending outside the dataset are possible as well as contacts from BCs with soma outside of the dataset, especially for larger types such as CBC8 and CBC9. The number of bulb contacts per CBC is underestimated as well since the true coverage factor of HC dendrites lies at about 5-7 while we have only five overlapping HCs in the center and coverage going down to one towards the edges of the dataset.

### Horizontal cell injections and immunolabeling for GABA receptors

Three HCs were injected using Alexa Fluor 568 as described before (Yadav, Tetenborg, and Dedek 2019). In brief, cell nuclei in the retinal whole-mount preparation were visualized with DAPI labeling. Based on depth and size of the nuclei, HCs were identified and then injected with Alexa Fluor 568 using sharp electrodes and subsequently fixed using 4% paraformaldehyde. Retinal whole-mounts were then incubated in primary antibodies, and immunolabeling for the GABA receptor subunit *γ*2 was carried as previously described (Ströh et al. 2013). Immunolabeling for VGAT and the GABA receptor subunit *γ*2 was carried out using fixed 12 μm thick vertical retina sections using standard protocols with primary antibodies against VGAT, the *γ*2 subunit and calbindin and secondary antibodies. All images were taken with a Leica TCS SP8 confocal microscope. Data was deconvolved with Huygens Essential software, using a theoretical point spread function and further processed using Fiji (Schindelin et al. 2012).

### Three-Dimensional Electron Microscopy using FIB-SEM

Focused ion beam-scanning electron microscopy (FIB-SEM tomography) allows efficient, complete, and automatic 3D reconstruction of HC dendrites with a resolution comparable to that of TEM (Xu et al. 2017; Bosch et al. 2015). An adult mouse (male, 14 weeks) was deeply anesthetized with isoflurane and decapitated before the eyes were dissected. All procedures were approved by the local animal care committee and were in accordance with the law for animal experiments issued by the German government (Tierschutzgesetz). The posterior eyecups were immersion fixed in a solution containing 0.1 M cacodylate buffer, 4% sucrose and 2% glutaraldehyde, and then rinsed in 0.15 M cacodylate buffer. A 1 × 1 mm^2^ retina piece was stained in a solution containing 1% osmium tetroxide, 1.5% potassium ferrocyanide, and 0.15 M cacodylate buffer. The osmium stain was amplified with 1% thiocarbohydrazide and 2% osmium tetroxide. The retina was then stained with 2% aqueous uranyl acetate and lead aspartate. The tissue was dehydrated through an 70-100% ethanol series, transferred to propylene oxide, infiltrated with 50%/50% propylene oxide/Epon Hard, and then 100% Epon Hard. The Epon Hard block was hardened at 60°C.

Afterwards, the block was prepared for FIB-SEM tomography. The sample was trimmed using an ultramicrotome (Leica UC 7) and afterwards glued onto a special sample stub (caesar workshop) using conductive silver paint. To avoid charge artifacts, all surfaces of the block were sputter-coated with 30 nm AuPd (80/20). A focused ion dual beam (FIB) workstation (XB 1540, Carl Zeiss Microscopy, Oberkochen, Germany) was used for tomogram acquisition. This instrument uses a focused gallium ion beam that can mill the sample at an angle of 54° with respect to the electron beam. A digital 24-bit scan-generator (ATLAS5, Carl Zeiss) was used to control ion and electron beam. The sample was milled using an ion beam of 1nA at an energy of 30 kV. Images were collected at an energy of 2 kV using a pixel size of 5 x 5 nm (x,y) and a layer thickness of 15 nm (z). Milling and imaging was performed simultaneously to compensate for charging effects. The raw images were converted into an image stack, black areas were cropped, and the images were aligned using cross correlation (Mastronarde 1997). HC dendrites were manually identified in ImageJ.

### Modelling

We built a biophysically realistic model of a HC dendritic branch using the simulation language NeuronC (Smith 1992). We used Vaa3D-Neuron2 auto-tracing (Xiao and Peng 2013) to generate a .swc file from the volume reconstruction of one HC branch and manually refined it in Neuromantic (Myatt et al. 2012). The model contains voltage-gated Ca^2+^ and K^+^ channels with different channel densities for proximal and distal dendrites and AMPA-type glutamate receptors at the cone synapses (Tab. 1). Photoreceptors were modelled as predefined in NeuronC with two compartments including voltage-gated Ca^2+^ and Ca^2+^-activated Cl^-^ channels. Cones were placed at the original positions with one synapse per invaginating HC dendritic tip found in the EM data. The synapses to the HC include postsynaptic AMPA channels modelled as Markov state-machines that included vesicle release noise. The model was stimulated for 60 s with both full-field and checkerboard Gaussian noise with a temporal frequency of 2 Hz and equal variance. The checkerboard stimulus consisted of independent pixels of 5 x 5 μm size such that all cones were stimulated independently. Voltage signals were recorded in a dendritic tip below each of the 12 cones and in 14 bulbs identified along the dendrite.

**Table 1.**
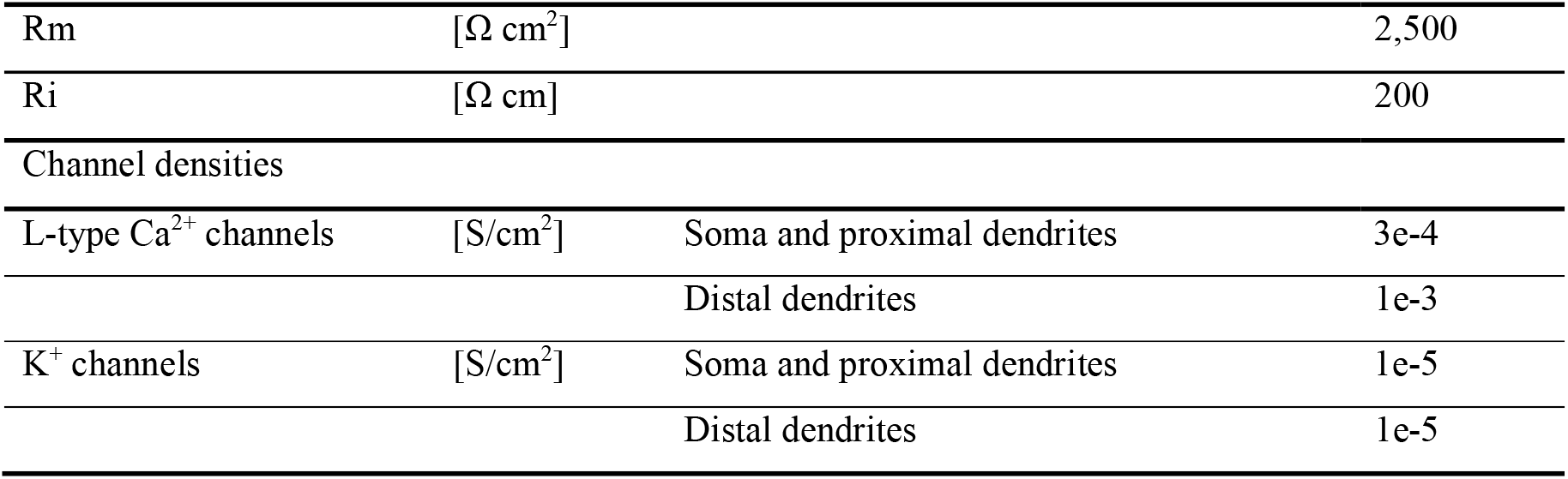
Parameters of the biophysical model.

### Statistics

Error bars in all plots are 95% confidence intervals (CIs) calculated as percentiles of the bootstrap distribution obtained via case resampling. In Figure 2c,d, we fitted generalized additive models (R package mgcv, Wood 2017) with Poisson output distribution for skeleton tips (Fig. 2c) and Gamma output distribution for contact area (Fig. 2d). Both had distance from soma as a smooth function and HC identity as smooth random effect.

## Acknowledgments

We thank Helmstaedter et al. 2013 for making their data available and L. Peichl for critical reading of the manuscript. This work was supported by the German Research Foundation (DFG) through individual grants (SCHU2243/3-1 to TS, BE5601/2-1 to PB) and the priority program SPP2041 (BE5601/4-1 to PB, EU 42/9-1 to TE), the German Ministry of Education and Research through the Bernstein Award (FKZ 01GQ1601 to PB) and NIH grants (EY023766 to TE & RGS; EY022070 to RGS).

## References

Barnes, S., Merchant, V., & Mahmud, F. 1993. “Modulation of Transmission Gain by Protons at the Photoreceptor Output Synapse.” Proceedings of the National Academy of Sciences 90 (21): 10081–85. https://doi.org/10.1073/pnas.90.21.10081.

Behrens, C., Schubert, T., Haverkamp, S., Euler, T., & Berens, P. 2016. “Connectivity Map of Bipolar Cells and Photoreceptors in the Mouse Retina.” ELife 5 (November): 065722. https://doi.org/10.7554/eLife.20041.

Borghuis, B.G., Ratliff, C.P., & Smith, R.G. 2018. “Impact of Light-Adaptive Mechanisms on Mammalian Retinal Visual Encoding at High Light Levels.” Journal of Neurophysiology 119 (4): 1437–49. https://doi.org/10.1152/jn.00682.2017.

Bosch, C., Martí-nez, A., Masachs, N., Teixeira, C.M., Fernaud, I., Ulloa, F., Pérez-Martí-nez, E., et al. 2015. “FIB/SEM Technology and High-Throughput 3D Reconstruction of Dendritic Spines and Synapses in GFP-Labeled Adult-Generated Neurons.” Frontiers in Neuroanatomy 9 (May). https://doi.org/10.3389/fnana.2015.00060.

Cardin, J.A. 2018. “Inhibitory Interneurons Regulate Temporal Precision and Correlations in Cortical Circuits.” Trends in Neurosciences 41 (10): 689–700. https://doi.org/10.1016/j.tins.2018.07.015.

Chapot, C.A., Behrens, C., Rogerson, L.E., Baden, T., Pop, S., Berens, P., Euler, T., & Schubert, T. 2017. “Local Signals in Mouse Horizontal Cell Dendrites.” Current Biology 27 (23): 3603–3615.e5. https://doi.org/10.1016/j.cub.2017.10.050.

Cueva, J.G., Haverkamp, S., Reimer, R.J., Edwards, R., Wässle, H., & Brecha, N.C. 2002. “Vesicular γ-Aminobutyric Acid Transporter Expression in Amacrine and Horizontal Cells.” Journal of Comparative Neurology 445 (3): 227–37. https://doi.org/10.1002/cne.10166.

Dedek, K., Breuninger, T., Sevilla Müller, L.P. de, Maxeiner, S., Schultz, K., Janssen-Bienhold, U., Willecke, K., Euler, T., & Weiler, R. 2009. “A Novel Type of Interplexiform Amacrine Cell in the Mouse Retina.” European Journal of Neuroscience 30 (2): 217–28. https://doi.org/10.1111/j.1460-9568.2009.06808.x.

Diamond, J.S. 2017. “Inhibitory Interneurons in the Retina: Types, Circuitry, and Function.” Annual Review of Vision Science 3 (1): 1–24. https://doi.org/10.1146/annurev-vision-102016-061345.

Dowling, J.E., & Boycott, B.B. 1966. “Organization of the Primate Retina: Electron Microscopy.” Proceedings of the Royal Society B: Biological Sciences 166 (1002): 80–111. https://doi.org/10.1098/rspb.1966.0086.

Dowling, J.E., Brown, J.E., & Major, D. 1966. “Synapses of Horizontal Cells in Rabbit and Cat Retinas.” Science 153 (3744): 1639–41. https://doi.org/10.1126/science.153.3744.1639.

Drinnenberg, A., Franke, F., Morikawa, R.K., Jüttner, J., Hillier, D., Hantz, P., Hierlemann, A., Azeredo da Silveira, R., & Roska, B. 2018. “How Diverse Retinal Functions Arise from Feedback at the First Visual Synapse.” Neuron 99 (1): 117–134.e11. https://doi.org/10.1016/j.neuron.2018.06.001.

Duebel, J., Haverkamp, S., Schleich, W., Feng, G., Augustine, G.J., Kuner, T., & Euler, T. 2006. “Two-Photon Imaging Reveals Somatodendritic Chloride Gradient in Retinal ON-Type Bipolar Cells Expressing the Biosensor Clomeleon.” Neuron 49 (1): 81–94. https://doi.org/10.1016/j.neuron.2005.10.035.

Ellias, S.A., & Stevens, J.K. 1980. “The Dendritic Varicosity: A Mechanism for Electrically Isolating the Dendrites of Cat Retinal Amacrine Cells?” Brain Research 196 (2): 365–72. https://doi.org/10.1016/0006-8993(80)90401-1.

Euler, T., Haverkamp, S., Schubert, T., & Baden, T. 2014. “Retinal Bipolar Cells: Elementary Building Blocks of Vision.” Nature Reviews Neuroscience 15 (8): 507–19. https://doi.org/10.1038/nrn3783.

Franke, K., Berens, P., Schubert, T., Bethge, M., Euler, T., & Baden, T. 2017. “Inhibition Decorrelates Visual Feature Representations in the Inner Retina.” Nature 542 (7642): 439–44. https://doi.org/10.1038/nature21394.

Gala, R., Lebrecht, D., Sahlender, D.A., Jorstad, A., Knott, G., Holtmaat, A., & Stepanyants, A. 2017. “Computer Assisted Detection of Axonal Bouton Structural Plasticity in in Vivo Time-Lapse Images.” ELife 6: 1–20. https://doi.org/10.7554/elife.29315.

Grove, J.C.R., Hirano, A.A., los Santos, J. de, McHugh, C.F., Purohit, S., Field, G.D., Brecha, N.C., & Barnes, S. 2019. “Novel Hybrid Action of GABA Mediates Inhibitory Feedback in the Mammalian Retina.” PLOS Biology 17 (4): e3000200. https://doi.org/10.1371/journal.pbio.3000200.

Guo, C., Stella, S.L., Hirano, A.A., & Brecha, N.C. 2009. “Plasmalemmal and Vesicular γ-Aminobutyric Acid Transporter Expression in the Developing Mouse Retina.” The Journal of Comparative Neurology 512 (1): 6–26. https://doi.org/10.1002/cne.21846.

Hartveit, E. 1999. “Reciprocal Synaptic Interactions Between Rod Bipolar Cells and Amacrine Cells in the Rat Retina.” Journal of Neurophysiology 81 (6): 2923–36. https://doi.org/10.1152/jn.1999.816.2923.

Haverkamp, S., Grünert, U., & Wässle, H. 2000. “The Cone Pedicle, a Complex Synapse in the Retina.” Neuron 27 (1): 85–95. https://doi.org/10.1016/S0896-6273(00)00011-8.

Haverkamp, S., & Wässle, H. 2000. “Immunocytochemical Analysis of the Mouse Retina.” The Journal of Comparative Neurology 424 (1): 1–23. https://doi.org/10.1002/1096-9861(20000814)424:1<1::AID-CNE1>3.0.CO;2-V.

He, S., Weiler, R., & Vaney, D.I. 2000. “Endogenous Dopaminergic Regulation of Horizontal Cell Coupling in the Mammalian Retina.” Journal of Comparative Neurology 418 (1): 33–40. https://doi.org/10.1002/(SICI)1096-9861(20000228)418:1<33::AID-CNE3>3.0.CO;2-J.

Helmstaedter, M., Briggman, K.L., & Denk, W. 2011. “High-Accuracy Neurite Reconstruction for High-Throughput Neuroanatomy.” Nature Neuroscience 14 (8): 1081–88. https://doi.org/10.1038/nn.2868.

Helmstaedter, M., Briggman, K.L., Turaga, S.C., Jain, V., Seung, H.S., & Denk, W. 2013. “Connectomic Reconstruction of the Inner Plexiform Layer in the Mouse Retina.” Nature 500 (7461): 168–74. https://doi.org/10.1038/nature12346.

Jackman, S.L., Babai, N., Chambers, J.J., Thoreson, W.B., & Kramer, R.H. 2011. “A Positive Feedback Synapse from Retinal Horizontal Cells to Cone Photoreceptors.” PLoS Biology 9 (5). https://doi.org/10.1371/journal.pbio.1001057.

Janssen-Bienhold, U., Trümpler, J., Hilgen, G., Schultz, K., Sevilla Muller, L.P. De, Sonntag, S., Dedek, K., Dirks, P., Willecke, K., & Weiler, R. 2009. “Connexin57 Is Expressed in Dendro-Dendritic and Axo-Axonal Gap Junctions of Mouse Horizontal Cells and Its Distribution Is Modulated by Light.” Journal of Comparative Neurology 513 (4): 363–74. https://doi.org/10.1002/cne.21965.

Kamermans, M., Fahrenfort, I., Schultz, K., Janssen-Bienhold, U., Sjoerdsma, T., & Weiler, R. 2001. “Hemichannel-Mediated Inhibition in the Outer Retina.” Science 292 (5519): 1178–80. https://doi.org/10.1126/science.1060101.

Kamermans, M., & Werblin, F.S. 1992. “GABA-Mediated Positive Autofeedback Loop Controls Horizontal Cell Kinetics in Tiger Salamander Retina.” The Journal of Neuroscience 12 (7): 2451–63. https://doi.org/10.1523/JNEUROSCI.12-07-02451.1992.

Kemmler, R., Schultz, K., Dedek, K., Euler, T., & Schubert, T. 2014. “Differential Regulation of Cone Calcium Signals by Different Horizontal Cell Feedback Mechanisms in the Mouse Retina.” Journal of Neuroscience 34 (35): 11826–43. https://doi.org/10.1523/JNEUROSCI.0272-14.2014.

Kolb, H. 1977. “The Organization of the Outer Plexiform Layer in the Retina of the Cat: Electron Microscopic Observations.” Journal of Neurocytology 6 (2): 131–53. https://doi.org/10.1007/BF01261502.

Lee, S., Zhang, Y., Chen, M., & Zhou, Z.J. 2016. “Segregated Glycine-Glutamate Co-Transmission from VGluT3 Amacrine Cells to Contrast-Suppressed and Contrast-Enhanced Retinal Circuits.” Neuron 90 (1): 27–34. https://doi.org/10.1016/j.neuron.2016.02.023.

Linberg, K.A., & Fisher, S.K. 1988. “Ultrastructural Evidence That Horizontal Cell Axon Terminals Are Presynaptic in the Human Retina.” The Journal of Comparative Neurology 268 (2): 281–97. https://doi.org/10.1002/cne.902680211.

Liu, X., Hirano, A.A., Sun, X., Brecha, N.C., & Barnes, S. 2013. “Calcium Channels in Rat Horizontal Cells Regulate Feedback Inhibition of Photoreceptors through an Unconventional GABA-and PH-Sensitive Mechanism.” The Journal of Physiology 591 (13): 3309–24. https://doi.org/10.1113/jphysiol.2012.248179.

Marchiafava, P.L. 1978. “Horizontal Cells Influence Membrane Potential of Bipolar Cells in the Retina of the Turtle.” Nature 275 (5676): 141–42. https://doi.org/10.1038/275141a0.

Mastronarde, D.N. 1997. “Dual-Axis Tomography: An Approach with Alignment Methods That Preserve Resolution.” Journal of Structural Biology 120 (3): 343–52. https://doi.org/10.1006/jsbi.1997.3919.

Miller, R.F., & Dacheux, R.F. 1976. “Synaptic Organization and Ionic Basis of on and off Channels in Mudpuppy Retina. I. Intracellular Analysis of Chloride-Sensitive Electrogenic Properties of Receptors, Horizontal Cells, Bipolar Cells, and Amacrine Cells.” The Journal of General Physiology 67 (6): 639–59. https://doi.org/10.1085/jgp.67.6.639.

Münch, T.A., Silveira, R.A. Da, Siegert, S., Viney, T.J., Awatramani, G.B., & Roska, B. 2009. “Approach Sensitivity in the Retina Processed by a Multifunctional Neural Circuit.” Nature Neuroscience 12 (10): 1308–16. https://doi.org/10.1038/nn.2389.

Myatt, D.R., Hadlington, T., Ascoli, G.A., & Nasuto, S.J. 2012. “Neuromantic – from Semi-Manual to Semi-Automatic Reconstruction of Neuron Morphology.” Frontiers in Neuroinformatics 6 (March): 1–14. https://doi.org/10.3389/fninf.2012.00004.

Olney, J.W. 1968. “An Electron Microscopic Study of Synapse Formation, Receptor Outer Segment Development, and Other Aspects of Developing Mouse Retina.” Investigative Ophthalmology & Visual Science 7 (3): 250–68. http://www.ncbi.nlm.nih.gov/pubmed/5655873.

Schindelin, J., Arganda-Carreras, I., Frise, E., Kaynig, V., Longair, M., Pietzsch, T., Preibisch, S., et al. 2012. “Fiji: An Open-Source Platform for Biological-Image Analysis.” Nature Methods 9 (7): 676–82. https://doi.org/10.1038/nmeth.2019.

Schubert, T., & Euler, T. 2010. “Retinal Processing: Global Players Like It Local.” Current Biology 20 (11): R486–88. https://doi.org/10.1016/j.cub.2010.04.034.

Schwartz, E.A. 1974. “Responses of Bipolar Cells in the Retina of the Turtle.” The Journal of Physiology 236 (1): 211–24. https://doi.org/10.1113/jphysiol.1974.sp010431.

Schwartz, E.A. 1987. “Depolarization without Calcium Can Release Gamma-Aminobutyric Acid from a Retinal Neuron.” Science 238 (4825): 350–55. https://doi.org/10.1126/science.2443977.

Smith, R.G. 1992. “NeuronC: A Computational Language for Investigating Functional Architecture of Neural Circuits.” Journal of Neuroscience Methods 43 (2-3): 83–108. https://doi.org/10.1016/0165-0270(92)90019-A.

Smith, R.G. 1995. “Simulation of an Anatomically Defined Local Circuit: The Cone-Horizontal Cell Network in Cat Retina.” Visual Neuroscience 12 (03): 545–61. https://doi.org/10.1017/S0952523800008440.

Srinivasan, M. V, Laughlin, S.B., & Dubs, A. 1982. “Predictive Coding: A Fresh View of Inhibition in the Retina.” Proceedings of the Royal Society of London. Series B. Biological Sciences 216 (1205): 427–59. https://doi.org/10.1098/rspb.1982.0085.

Ströh, S., Puller, C., Swirski, S., Hölzel, M.-B., Linde, L.I.S. van der, Segelken, J., Schultz, K., et al. 2018. “Eliminating Glutamatergic Input onto Horizontal Cells Changes the Dynamic Range and Receptive Field Organization of Mouse Retinal Ganglion Cells.” The Journal of Neuroscience 38 (8): 0141–17. https://doi.org/10.1523/JNEUROSCI.0141-17.2018.

Ströh, S., Sonntag, S., Janssen-Bienhold, U., Schultz, K., Cimiotti, K., Weiler, R., Willecke, K., & Dedek, K. 2013. “Cell-Specific Cre Recombinase Expression Allows Selective Ablation of Glutamate Receptors from Mouse Horizontal Cells.” PLoS ONE 8 (12): 15–17. https://doi.org/10.1371/journal.pone.0083076.

Taylor, W.R., & Smith, R.G. 2012. “The Role of Starburst Amacrine Cells in Visual Signal Processing.” Visual Neuroscience 29 (01): 73–81. https://doi.org/10.1017/S0952523811000393.

Thoreson, W.B., & Mangel, S.C. 2012. “Lateral Interactions in the Outer Retina.” Progress in Retinal and Eye Research 31 (5): 407–41. https://doi.org/10.1016/j.preteyeres.2012.04.003.

Tien, N.W., Kim, T., & Kerschensteiner, D. 2016. “Target-Specific Glycinergic Transmission from VGluT3-Expressing Amacrine Cells Shapes Suppressive Contrast Responses in the Retina.” Cell Reports 15 (7): 1369–75. https://doi.org/10.1016/j.celrep.2016.04.025.

Tsukamoto, Y., & Omi, N. 2014. “Some OFF Bipolar Cell Types Make Contact with Both Rods and Cones in Macaque and Mouse Retinas.” Frontiers in Neuroanatomy 8 (September): 105. https://doi.org/10.3389/fnana.2014.00105.

Turrigiano, G.G. 2011. “Stabilizing Neuronal Function Homeostatic Synaptic Plasticity: Local and Global Mechanisms for Homeostatic Synaptic Plasticity: Local and Global Mechanisms for Stabilizing Neuronal Function.” Cold Spring Harb Perspect Biol, 1–18. https://doi.org/10.1101/cshperspect.a005736.

Vardi, N., Zhang, L.-L., Payne, J.A., & Sterling, P. 2000. “Evidence That Different Cation Chloride Cotransporters in Retinal Neurons Allow Opposite Responses to GABA.” The Journal of Neuroscience 20 (20): 7657–63. https://doi.org/10.1523/JNEUROSCI.20-20-07657.2000.

Verweij, J., Kamermans, M., & Spekreijse, H. 1996. “Horizontal Cells Feed Back to Cones by Shifting the Cone Calcium-Current Activation Range.” Vision Research 36 (24): 3943–53. https://doi.org/10.1016/S0042-6989(96)00142-3.

Vroman, R., Klaassen, L.J., Howlett, M.H.C., Cenedese, V., Klooster, J., Sjoerdsma, T., & Kamermans, M. 2014. “Extracellular ATP Hydrolysis Inhibits Synaptic Transmission by Increasing PH Buffering in the Synaptic Cleft.” PLoS Biology 12 (5). https://doi.org/10.1371/journal.pbio.1001864.

Warren, T.J., Hook, M.J. Van, Supuran, C.T., & Thoreson, W.B. 2016. “Sources of Protons and a Role for Bicarbonate in Inhibitory Feedback from Horizontal Cells to Cones in Ambystoma Tigrinum Retina.” Journal of Physiology 594 (22): 6661–77. https://doi.org/10.1113/JP272533.

Wässle, H., Puller, C., Müller, F., & Haverkamp, S. 2009. “Cone Contacts, Mosaics, and Territories of Bipolar Cells in the Mouse Retina.” The Journal of Neuroscience: The Official Journal of the Society for Neuroscience 29 (1): 106–17. https://doi.org/10.1523/JNEUROSCI.4442-08.2009.

Werblin, F.S., & Dowling, J.E. 1969. “Organization of the Retina of the Mudpuppy, Necturus Maculosus. II. Intracellular Recording.” Journal of Neurophysiology 32 (3): 339–55. https://doi.org/10.1152/jn.1969.32.3.339.

Witkovsky, P., Gábriel, R., & Križaj, D. 2008. “Anatomical and Neurochemical Characterization of Dopaminergic Interplexiform Processes in Mouse and Rat Retinas.” The Journal of Comparative Neurology 510 (2): 158–74. https://doi.org/10.1002/cne.21784.

Wood, S.N. 2017. Generalized Additive Models: An Introduction with R, Second Edition. 2nd ed. Chapman and Hall/CRC.

Xiao, H., & Peng, H. 2013. “APP2: Automatic Tracing of 3D Neuron Morphology Based on Hierarchical Pruning of a Gray-Weighted Image Distance-Tree.” Bioinformatics 29 (11): 1448–54. https://doi.org/10.1093/bioinformatics/btt170.

Xu, C.S., Hayworth, K.J., Lu, Z., Grob, P., Hassan, A.M., García-Cerdán, J.G., Niyogi, K.K., Nogales, E., Weinberg, R.J., & Hess, H.F. 2017. “Enhanced FIB-SEM Systems for Large-Volume 3D Imaging.” ELife 6: 1–36. https://doi.org/10.7554/eLife.25916.

Yadav, S.C., Tetenborg, S., & Dedek, K. 2019. “Corrigendum: Gap Junctions in A8 Amacrine Cells Are Made of Connexin36 but Are Differently Regulated Than Gap Junctions in AII Amacrine Cells.” Frontiers in Molecular Neuroscience 12 (June): 1–2. https://doi.org/10.3389/fnmol.2019.00149.

Yang, X., & Wu, S. 1991. “Feedforward Lateral Inhibition in Retinal Bipolar Cells: Input-Output Relation of the Horizontal Cell-Depolarizing Bipolar Cell Synapse.” Pnas 88 (8): 3310–13. https://doi.org/10.1073/pnas.88.8.3310.

